# Linking Transcriptional and Cellular Responses of Human and Murine Pneumococcal Pneumonia

**DOI:** 10.1101/2025.02.13.638069

**Authors:** Anna-Dorothea Gorki, Karin Lakovits, Anastasiya Hladik, Ildiko Mesteri, Riem Gawish, Rui Martins, Barbara Drobits, Jörg Menche, Sylvia Knapp, Stefanie Widder

## Abstract

Community-acquired pneumonia (CAP), often caused by *Streptococcus pneumoniae*, poses a significant global health challenge, especially among high-risk populations. This study investigates the temporal dynamics of pulmonary gene expression during *S. pneumoniae*-induced pneumonia using a murine model to elucidate host-pathogen interactions and identify potential biomarkers of disease severity. Using bulk RNA sequencing, we analyzed lung tissues at early (1h, 8h) and acute infection (2d, 3d), as well as post-resolution (30d) time-points. At 2 days post-infection, differentially expressed genes (DEGs) revealed heightened innate immune responses, including chemokine and interferon signaling pathways. By 30 days post-infection, gene expression profiles and histological changes normalized, reflecting resolution of inflammation. Stratifying mice into recovered, sick, and moribund phenotypes during the acute infection phase highlighted significant transcriptional distinctions, including upregulated pro-inflammatory genes and Schlafen family members. Arginase 1 emerged as a predictive marker of disease severity, with elevated expression detectable as early as 8 hours post-infection. Comparative analyses revealed significant overlap between transcriptional responses in our model and those from a Gram-negative *Acinetobacter baumannii* pneumonia model, implicating conserved pathways, such as IL-17 and Toll-like receptor signaling. Additionally, murine DEGs correlated with human plasma proteins associated with severe CAP, including CCL8 and CD14, suggesting translational relevance.

This study underscores phase-specific transcriptional reprogramming during pneumococcal pneumonia and identifies potential biomarkers and therapeutic targets for improving outcomes in severe CAP cases.

## Introduction

Community-acquired pneumonia (CAP) remains a leading cause of morbidity and mortality worldwide, with particularly high fatality rates among patients requiring intensive care, where mortality can reach up to 50% (1). *Streptococcus pneumoniae* (*S. pneumoniae*), a Gram-positive bacterium, is the most common pathogen responsible for CAP (2). While *S. pneumoniae* is commonly carried asymptomatically in the nasopharynx, it can migrate to the lungs, leading to serious diseases like pneumonia especially in young children, elderly and immunocompromised individuals (3,4). Despite advances in vaccine development, antimicrobial therapies and supportive care, the annual death rate caused by pneumonia especially in elderly is still alarmingly high (5). This emphasizes the need for deeper insights into the host-pathogen interactions that drive pneumonia outcomes. The progression of bacterial pneumonia involves complex changes in gene expression and reorganization of the pulmonary architecture that mediate both acute host responses and long-term recovery. During infection, initial bacterial colonization can escalate into severe lung damage, which often extends to systemic bacterial dissemination. Murine models of bacterial pneumonia offer a powerful tool for dissecting the temporal dynamics of host responses to *S. pneumoniae* infection. Previous works have shown the importance of e.g. alveolar macrophages (6,7) and neutrophils (8–10) during pneumococcal pneumonia on a cellular level and type I interferon (11) and NF-κB signaling (12) on a molecular level. However, temporal dynamics of the disease course and the changes of lung gene expression throughout are less well studied.

Here we studied the longitudinal progression of *S. pneumoniae* infection using a murine model. By analyzing pulmonary transcriptional profiles across early and acute infection time-points as well as a post-resolution time-point, we identified critical pathways involved in both the immediate immune response and subsequent tissue repair mechanisms. Moreover, the comparison with previously published data of a murine pneumonia model induced by the Gram-negative bacterium *Acinetobacter baumannii* (13) uncovered a conserved immune response across both pneumonia models. Most importantly, we found differentially expressed genes in the model that aligned with human plasma proteins and correlated with pneumonia severity (14), suggesting their potential as markers of severe pneumonia. Our analysis further revealed distinct transcriptional shifts between sickness states, providing new insights into the molecular drivers of pneumonia. These findings not only enhance our understanding of the disease but also highlights potential therapeutic targets that could improve outcomes for patients with pneumococcal pneumonia.

## Material and Methods

### Mouse Pneumonia Model and Sampling

Female wildtype C57BL/6J mice, 8 to 9 weeks old were purchased from Charles River and housed under specific pathogen-free conditions at the Medical University of Vienna for at least 1 week prior to experiment. Mice were infected intranasally with 1× 10^4^ CFUs *S. pneumoniae* serotype 3 (ATCC 6303) as described before (15,16). Every 6 to 8 hours phenotypic assessment was performed. Animal experiments were approved by the Austrian Federal Ministry of Sciences and Research (BMWF-66.009/0285-II/3b/2011). Whole lungs, spleen, liver and blood were harvested at indicated times after inoculation with *S. pneumoniae* or sterile saline (controls). Colony-forming units (CFUs) in lung, liver, spleen and blood were determined by 10-fold serial dilutions of homogenates on blood agar plates.

### RNA Extraction, Library Construction and Sequencing

Lung samples for RNA sequencing were homogenized in 0.5mL of TRIzol reagent (Invitrogen) and transferred onto QiaShredder (Qiagen). Total cellular RNA was isolated from lung tissues by Maxwell® extraction according to the manufacturer’s protocol and stored at −80°C. Afterwards, RNA was quantified and its quality was assessed using the Qubit 2.0 Fluorometric Quantitation system (Life Technologies) and the Experion Automated Electrophoresis System (Bio-Rad). Libraries were sequenced at the Biomedical Sequencing Facility at the CeMM Research Center for Molecular Medicine of the Austrian Academy of Sciences using the HiSeq 3000/4000 platform (Illumina) in a 25bp single-read configuration. Data was processed as previously described (17).

### Differential Gene Expression and Pathway Analysis

Differential gene expression was performed using DeSeq2 (18). Genes with low expression (base mean count ≤50) were excluded, and thresholds of absolute log2 fold-change >2 and adjusted p-value <0.05 were applied. GO term and KEGG Pathway analyses were performed using the ShinyGO 0.81 platform (19). Venn diagrams preparation was supported by following tool: https://bioinformatics.psb.ugent.be/webtools/Venn/.

### Immune cell prediction

The ImmuCellAI-mouse tool (20) was used, which employs a hierarchical classification strategy with three layers to predict immune cell composition in RNA Seq samples. Layer 1 consists of three types of cells of the lymphoid lineage (B-cells, NK cells, and T-cells) and four types of the myeloid lineage (macrophages, dendritic cells, monocytes, and granulocytes). Layer 2 includes subtypes of the cells found in layer 1, while layer 3 comprises additional subtypes of CD4 T-cells and CD8 T-cells (layer 2).

### Histopathology Analysis

Paraffin-embedded lung sections were stained with H&E and scored by a trained pathologist who was blinded to experimental groups. The final pneumonia score was the sum of the following parameters: severity of pleuritis, interstitial inflammation, edema, and pneumonia were scored as 0= absent, 1= mild, 2= moderately severe, 3= severe; bronchitis and pneumonia were scored as 1 if present; endothelitis was scored as 0= absent, 2= present, 3= present with endothelial wall necrosis and a score of 0.5 was added for every infiltrate covering 10% of the lung area.

### Statistical Analysis

All statistical analyses were performed using Graph Pad Prism software version 10 or R platform (21). The significance between groups was statistically evaluated using a one-way or two-way ANOVA test followed by Sidak’s multiple comparison test for all figures except for immune cell prediction (Fig.3A-D, FigS3) were a Wilcoxon signed-rank test was performed. A p<0.05 was regarded as statistically significant, with *p<0.05, **p<0.005 and ***p<0.001.

## Results

### Acute and long-term alterations of pulmonary gene expression after pneumonia infection

To study how pulmonary gene expression profiles change during the course of bacterial pneumonia, we analyzed early (1h, 8h), acute (2d, 3d), and post-resolution (30d) time-points in a murine model of *S. pneumoniae* infection (Figure S1A). Mice were challenged with a dose of 1×10^4^ CFU of *S. pneumoniae*, corresponding to an LD50 dose in our set-up. Mortality occurred between 42- and 96 hours post-infection, resulting in 55% survival of infected mice until the end of the observation interval (Figure S1B). We collected lung, spleen, liver, and blood samples from both infected and control mice and first analyzed an acute infection (2dpi) and post-resolution (30dpi) time-point to identify changes associated with both phases. At 2dpi, we detected a significant bacterial load in the lungs of infected mice, accompanied by a significant reduction in body weight compared to baseline (Figure 1A). Systemic bacterial dissemination was confirmed by the presence of bacteria in the blood (Figure 1B), spleen, and liver (Figure S1C). Histological analysis of lung tissue revealed significantly elevated pneumonia scores at 2dpi due to edema formation, bronchitis, interstitial inflammation and cellular infiltrates (Figure 1C, Figure S1D). At post-resolution time-point (30dpi), the severity of the histological changes decreased as pneumonia was fully resolved and there was no longer a statistically significant difference between infected and control mice detectable, only interstitial inflammation remained slightly elevated (Figure 1C, Figure S1D).

**Figure 1.**
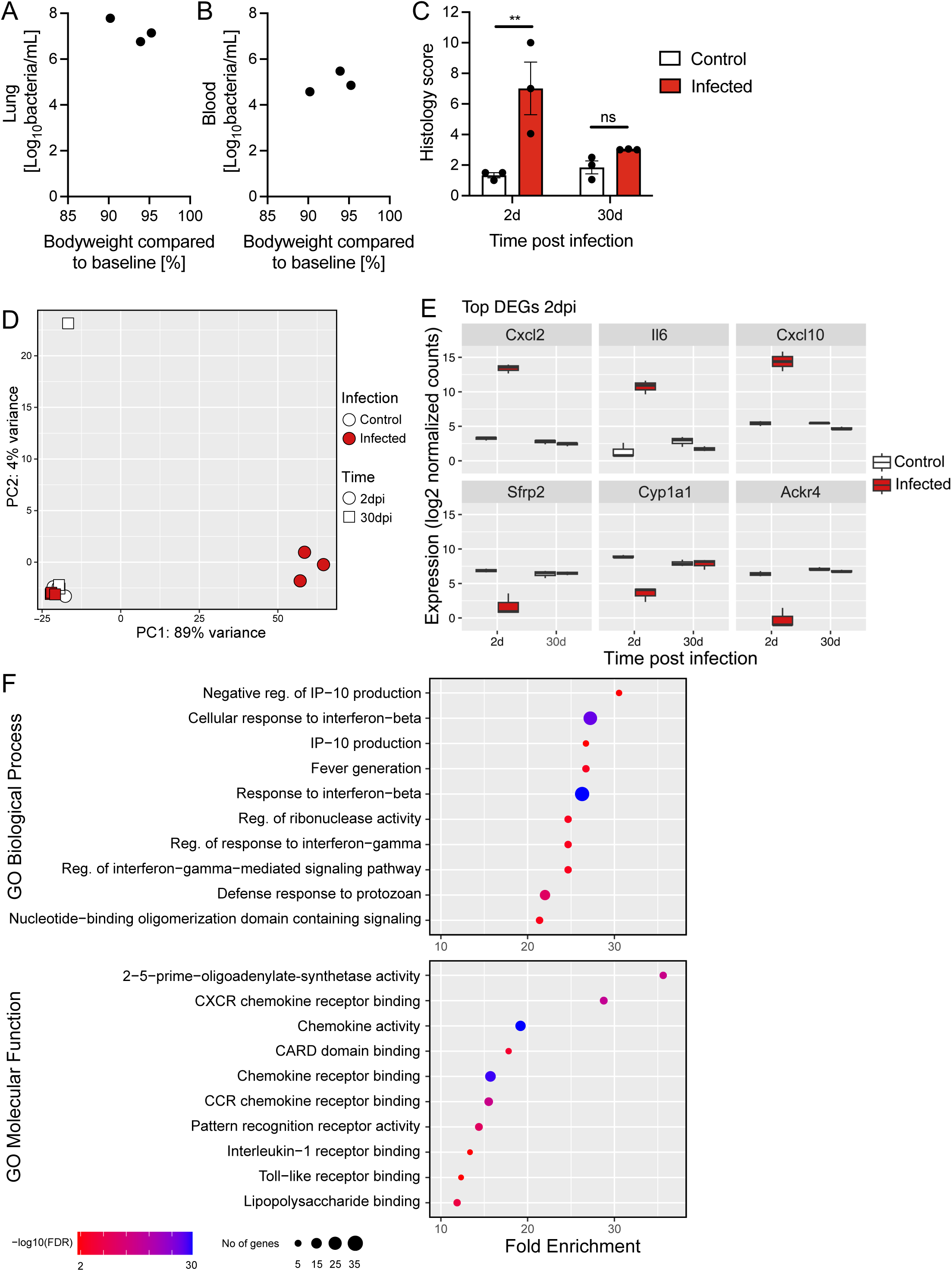
Acute and long-term alterations of pulmonary gene expression after pneumonia infection. Experimental setup of study used for (**A to F**): Female C57BL/6J WT mice were infected with 1×10^4^ CFU *S.pneumoniae* (infected) or given saline (control). Organs and blood were taken at indicated times post-infection. (**A to B**) Bacterial count in lung (**A**) and blood (**B**) 2 days post-infection (2dpi) versus bodyweight compared to baseline. (**C**) Lung histology score in infected and control mice at indicated time-points post-infection. (**D**) PCA of transcriptomic lung data of infected and control mice 2dpi and 30dpi. (**E**) Top three up- and downregulated DEGs (absolute log2fc >2, padj <0.05, base mean count >50) 2dpi based on log2 fold change in infected versus control mice. (**F**) Pathway analysis of DEGs in infected versus control mice 2dpi. CFU= Colony forming unit, DEG= Differentially expressed genes, Dpi= Days post infection, fc=fold change, GO=Gene ontology, PCA=Principal component analysis, WT=wild-type

We performed bulk RNA sequencing on lung tissue and using principal component analysis (PCA) and sample distance matrix analysis, we observed pronounced transcriptional differences in the acute and post-resolution phase (Figure 1D, Figure S1E). Our analysis revealed two independent clusters: one consisting of samples from infected mice at the acute infection (2dpi), and a second cluster containing all control samples and samples from the post-resolution phase (30dpi). To identify the most important expression characteristics of both phases, we performed a differential gene expression analysis for each time-point. We identified 316 differentially expressed genes (DEGs) at 2dpi compared to controls, with 299 genes qualifying as upregulated and 17 as downregulated (Table S1). By contrast, in total only two downregulated DEGs were detected at 30dpi compared to controls (Figure S1F, Table S2). Among the top upregulated genes which displayed the highest fold changes at 2dpi were pro-inflammatory mediators such as C-X-C motif chemokine ligand 2 (*Cxcl2*), Interleukin 6 (*Il6*), and *Cxcl10* (Figure 1E, upper panel). The most significantly downregulated genes were atypical chemokine receptor 4 (*Ackr4*), cytochrome P450, family 1, subfamily a, polypeptide 1 (*Cyp1a1*), and secreted frizzled-related protein 2 (*Sfrp2*) (Figure 1E, lower panel). Next, we conducted a pathway enrichment analysis to specify molecular processes associated with acute pneumonia (2dpi). We detected significant enrichment in interferon signaling (Figure 1F, GO Biological Process), complemented by pathways for chemokine and chemokine receptor signaling as well as Toll-like receptor binding and interleukin-1 receptor binding (Figure 1F, GO Molecular Function). This result underscores the crucial role of innate defense mechanisms in acute pneumococcal pneumonia.

### Classification of distinct sickness states during acute infection phase

At 2- and 3 days post infection, the acute infection phase, *S. pneumoniae*-infected mice exhibited distinct behavioral and physical differences. While some mice remained active with shiny fur, others displayed slightly sluggish reactions, and some had ruffled fur, reduced body temperature compared to uninfected controls, as well as severely limited reactions. Based on these observations, we combined the mice for the acute infection phase (2- and 3dpi) and classified them into three sickness states: recovered, sick, and moribund. This phenotypic classification was supported by the ordination of the lung transcriptomes (Figure 2A, Figure S2A), displaying a clear separation among sickness states. In detail, moribund mice exhibited the highest bacterial burdens in the lung, spleen, and liver, along with significant body weight loss of up to 13% compared to baseline (Figure 2B, Figures S2B-C). Sick mice displayed intermediate bacterial burdens and body weight loss (Figure 2B, Figures S2B-C), and recovered mice had negligible or undetectable bacterial loads and only a slight reduction in body weight. Furthermore, bacteria were detected in the blood of all sick and moribund mice, with no clear distinction in bacterial numbers between these groups, consistent with systemic infection (Figure 2C). Recovered mice, however, had minimal or undetectable bacterial presence in the blood (Figure 2C). Gene expression analysis of the sickness states revealed no DEGs in recovered mice compared to uninfected controls. In contrast, we identified 479 DEGs in the sick state (Table S3) and 783 DEGs in the moribund state (Table S4) compared to uninfected controls, with the majority being upregulated: 468 DEGs in the sick state and 631 DEGs in the moribund state. Shared between the sick and moribund groups were 438 upregulated DEGs (Figure 2D) alongside 8 downregulated DEGs (Figure 2E). These shared transcriptional changes represent the core host response to *S. pneumoniae* infection in our model. Notable upregulated genes included well-characterized drivers of the pneumonia response, such as Toll-like receptors *Tlr2* (22), *Tlr13* (23), *Tlr6*, as well as *Myd88* (24), *Nod2* (25), *Lcn2* (16), *Il10* (26), *Cxcl1* (27), and *Cxcl2* (28). Additionally, several members of the Schlafen family (*Slfn1, 2, 3, 4, 5, 8, 9, 10*) were upregulated, although their roles in pneumonia are not well understood. In summary, *S. pneumoniae*-infected mice at the peak of infection display distinct sickness states—recovered, sick, and moribund—that are characterized by differences in behavior, appearance, bacterial load, body weight loss, and pulmonary gene expression levels. Moribund mice exhibited the highest bacterial burden and most pronounced transcriptional alterations.

**Figure 2.**
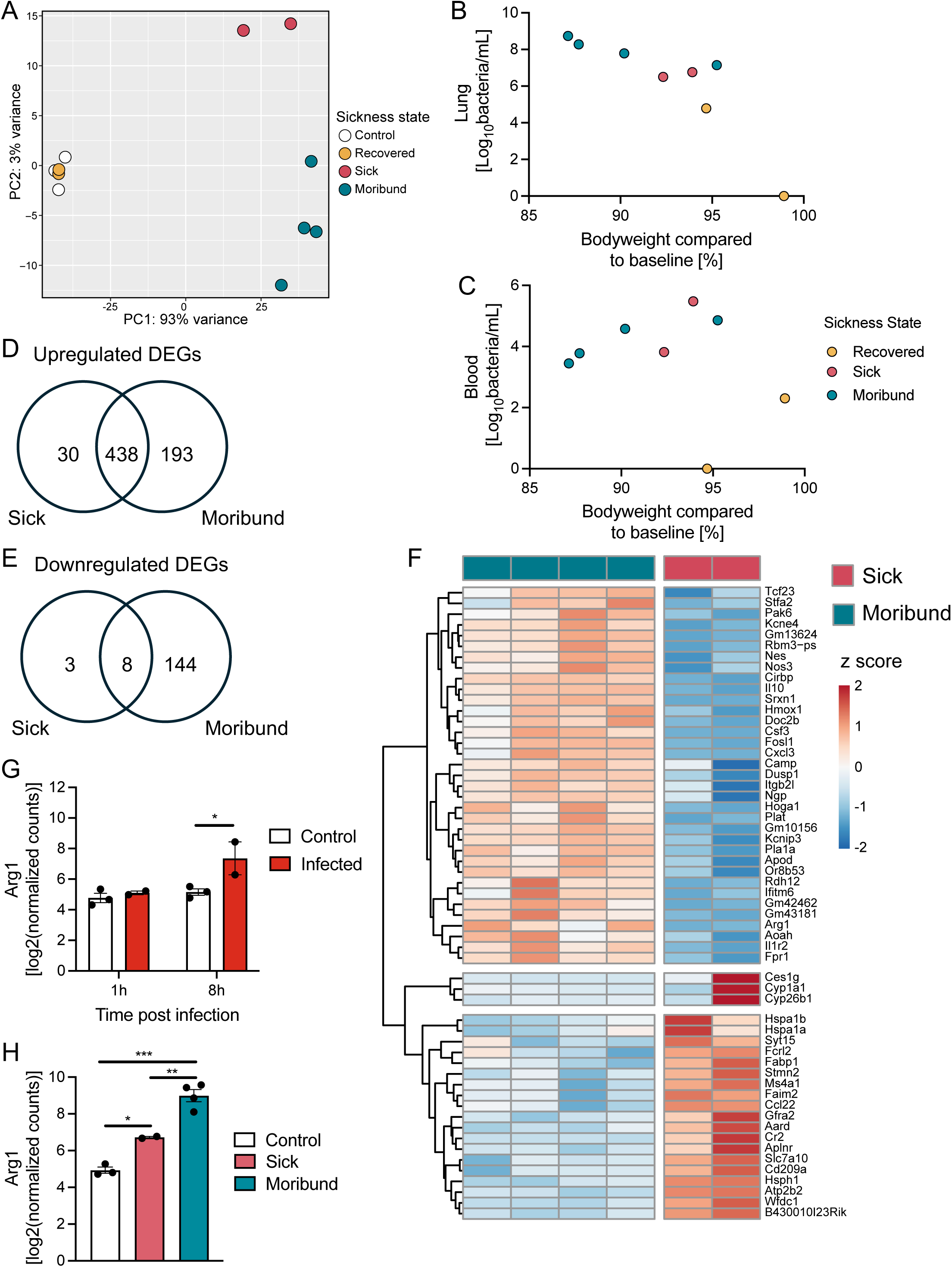
Classification of distinct sickness states during acute infection phase. Three different sickness stages (recovered, sick, moribund) were observed in *S. pneumoniae* infected mice during acute infection phase (2- and 3dpi). (**A**) PCA of RNA seq data of infected mice (recovered, sick, moribund) and controls. (**B and C**) Bacterial count in lung (**B**) and blood (**C**) of infected mice (recovered, sick, moribund) versus bodyweight compared to baseline. (**D to E**) Venn diagram of upregulated (**D**) and downregulated (**E**) DEGs (absolute log2fc >2, padj <0.05, base mean count >50) in sick and moribund mice compared to uninfected controls. (**F**) Heatmap of 57 DEGs (absolute log2fc >2, padj <0.05) between sick and moribund mice. (**G and H**) Arginase 1 (Arg1) expression at 1h and 8hpi (**G**) and in control, sick and moribund mice (**H**). Dpi= days post infection

### Identification of early diagnostic markers for moribund state in pneumonia

To assess, whether early differences in gene expression could predict future disease trajectories, we searched for significant differences between the sick and moribund state in the acute infection phase (2- and 3dpi) and compared the resulting DEGs to early infection time-points. Differential gene expression analysis between sick and moribund mice resulted in 57 DEGs. Using hierarchical clustering we identified three distinct gene clusters (Figure 2F). Cluster 1 included genes with higher expression in moribund mice, including p21 (RAC1) activated kinase 6 (*Pak6*), which was shown to be a region of interest for single-nucleotide polymorphism in childhood pneumonia (29) and interleukin-10 (*Il10*), which has been linked to the severity of systemic inflammatory response syndrome in community-acquired pneumonia (30). Cluster 2 comprised three genes influenced by inconsistency between samples in the sick group; these genes were excluded from further analysis. Cluster 3 contained genes more highly expressed in sick mice, including heat shock proteins (*Hspa1a*, *Hspa1b*, *Hsph1*) and immune-related genes (*Ccl22*, *Cd209a*).

Next, we investigated whether the expression of genes that differentiate between sickness states of sick and moribund was already altered at early infection time-points (1- and 8hpi) and might serve as predictor for disease severity. While bacterial presence in the lungs was confirmed at both time-points (Figure S2D), no DEGs were identified at 1hpi compared to uninfected controls. In contrast, 29 DEGs were detected at 8hpi (Table S5). These included early expression of Arginase 1 (*Arg1*), a gene that also distinguished moribund and sick mice during the acute infection phase. Expression analysis of *Arg1* showed significantly elevated levels in infected mice at 8hpi compared to uninfected controls (Figure 2G). This increase was also observed in sick mice at the peak of infection, with moribund mice exhibiting markedly higher *Arg1* expression compared to both uninfected and sick mice (Figure 2H). These findings are consistent with reports indicating that elevated Arginase 1 activity impairs lung-protective immunity during *S. pneumoniae* infection (31).

### Immune cell dynamics in moribund mice during acute pneumonia

Complementary to the gene expression analysis, we modeled the cellular immune response in *S. pneumoniae*-infected moribund mice at the acute infection phase (2- and 3dpi) using the ImmuCellAI-mouse tool (20). This tool estimates the likelihood of cellular infiltration across 36 immune cell types. The moribund state was characterized by a significant reduction in B-cells compared to uninfected controls, with trends indicating reduced dendritic cells, NK cells, and T-cells (Figure 3A, layer 1). Cell subtype analysis (layer 2) showed that these changes were driven primarily by a loss of B1- and plasma cells (Figure 3B), plasmacytoid dendritic cells (pDCs) (Figure S3A), NKT and CD4 T-cells (Figure S3B). Conversely, the model predicted increased macrophages, monocytes, and granulocytes in moribund mice. Further analysis of granulocyte subtypes identified increased neutrophils and reduced eosinophils in moribund mice (Figure 3C). Among macrophages, M1 like macrophages were significantly elevated, while M2 like macrophages remained unchanged compared to uninfected controls (Figure 3D). These findings align with prior research highlighting the role of alveolar macrophages, inflammatory monocytes, and neutrophil influx during pneumonia (6,10,32) but also highlighting the importance of B1 cells in the acute infection phase (33).

**Figure 3.**
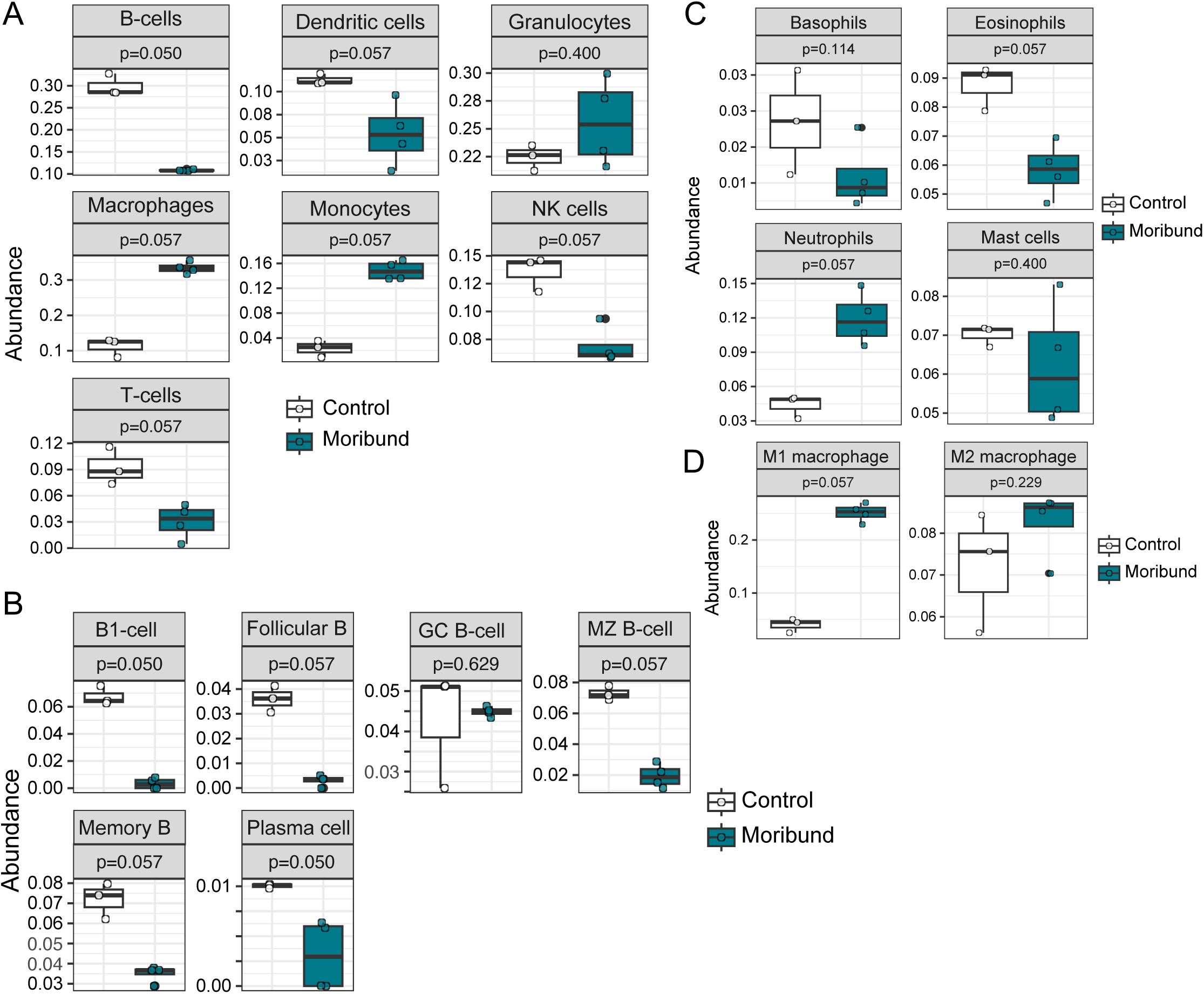
Immune cell dynamics in moribund mice during acute pneumonia. (**A to D**) Cell infiltration in lung tissue based on ImmuCellAI calculation for layer 1 cell types (**A**) and layer 2 B-cell (**B**), granulocyte (**C**) and macrophage (**D**) subtypes based on transcriptomic data of moribund mice and controls during acute infection phase (2- and 3dpi).

### Communal pulmonary gene expression patterns in murine and human pneumonia responses

Next, we examined whether we detect similarities in response patterns across pneumonia models caused by different pathogens. We compared DEGs from moribund mice at the acute infection phase (2- and 3dpi) to those identified in an acute lethal pneumonia model caused by *Acinetobacter baumannii*, a Gram-negative pathogen associated with hospital-acquired infections (13). The study by Zeng et al. focused on innate immune responses using Rag1 knockout mice deficient in functional T and B cells and analyzed lung tissue RNA at the acute phase of infection (24 hours post-infection). Even with differences in the utilized mouse model, we identified 472 overlapping upregulated genes and 74 overlapping downregulated genes between the two models (Figure 4A-B, Table S6). KEGG pathway analysis of the 546 shared DEGs revealed enrichment of pathways including IL-17, TNF, Toll-like receptor, and NOD-like receptor signaling (Figure 4C, Table S7), highlighting a shared innate immune response to severe bacterial pneumonia.

**Figure 4.**
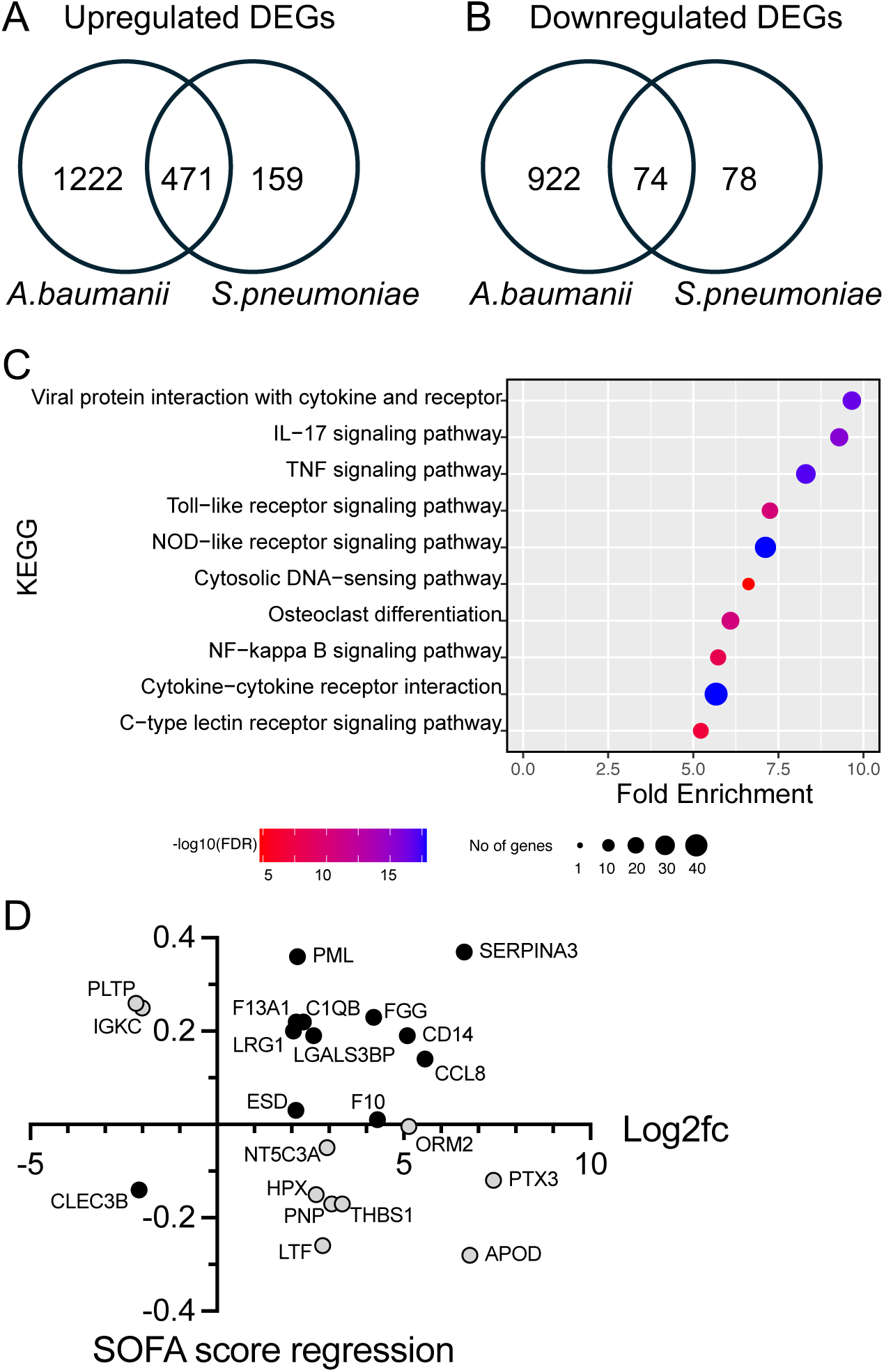
Communal pulmonary gene expression patterns in murine and human pneumonia responses. (**A and B**) Overview of up- and downregulated DEGs (absolute log2fc >2, padj <0.05, base mean count >50) in moribund mice infected by *S. pneumoniae* during acute infection phase (2- and 3dpi) in comparison to published lung mouse dataset using an *A. baumanii* induced acute lethal pneumonia model (absolute log2fc >2, p-value <0.01). (**C**) Enriched KEGG pathways of overlapping DEGs between *S. pneumoniae* and *A. baumanii* infection. (**D**) Moribund DEGs (absolute log2fc >2, padj <0.05, base mean count >50) were converted to human symbols and compared to published dataset on SOFA score correlation to plasma proteomics data in CAP patients. CAP= Community-acquired pneumonia

Finally, we assessed whether DEGs from moribund mice at the acute infection phase (2- and 3dpi) could serve as markers for severe pneumonia in human patients. The Sequential Organ Failure Assessment (SOFA) score is an approach to evaluate and track organ dysfunction in critically ill patients and therefore an effective tool for identifying severe CAP cases (34). We compared our murine DEGs to a published dataset of human plasma proteins associated with the SOFA score in community-acquired pneumonia (CAP) patients (14). Among the plasma proteins associated with SOFA score, 11 proteins exhibited a positive correlation with SOFA and were upregulated on gene level in moribund mice, including C-C motif chemokine 8 (*CCL8*), complement C1q (*C1QB*), and monocyte differentiation antigen CD14 (*CD14*) (Figure 4F, upper right quadrant). Tetranectin (*CLEC3B*), was the only plasma protein that expressed a negative association to SOFA as well as a downregulation on gene level in moribund mice (Figure 4F, lower left quadrant). For the remaining overlapping plasma proteins, association tendency to SOFA score differed to our murine lung RNA expression results (Figure 4F, upper left and lower right quadrant, grey dots).

Overall, our analyses identified a communal response to Gram-positive and Gram-negative induced pneumonia with shared pulmonary expression patterns and immune pathways (IL-17 and Toll-like receptor pathways) in moribund mice. Moreover, acute murine gene-level changes aligned with human plasma proteins detected in severe CAP patients and associated via the clinical SOFA score.

## Discussion

Our study elucidates the specific changes in pulmonary gene expression during bacterial pneumonia and revealed highly orchestrated, phase-specific responses involving significant transcriptional reprogramming during acute infection in a murine model of *S. pneumoniae* infection. Pathological changes at the peak of infection correlated with extensive transcriptional remodeling. This included the increase of pro-inflammatory mediators such as *Cxcl2*, *Il6*, and *Cxcl10*, alongside pathways associated with interferon signaling and chemokine activity. The expression changes indicated the activation of innate immune defenses aimed at combating the bacterial burden and aligned with previous reports (35,36). Interestingly, several novel drivers of the acute inflammatory response were detected, including aconitate decarboxylase-1 (*Acod1*, also known as Irg1), T cell-interacting, activating receptor on myeloid cells 1 (*Tarm1*) and oligoadenylate synthase (Oas) genes (*Oasl2, Oas3, Oas2, Oas1a, Oas1g, Oasl1*), all with unknown roles in pneumococcal pneumonia. The simultaneous downregulation of genes like Ackr4 and Cyp1a1, which are associated with anti-inflammatory or metabolic pathways (37,38), suggested a trade-off favoring host defense over tissue homeostasis during acute infection. By 30dpi, the pulmonary transcriptome had largely returned to baseline, reflecting resolution of inflammation and recovery. The absence of significant histological differences and the minimal number of DEGs at this stage demonstrated the capacity of the host to restore homeostasis following the resolution of infection.

A nuanced analysis of phenotypes at the acute infection phase (2- and 3dpi) stratified disease progression into three states: sick, moribund and recovered. The moribund state exhibited the most pronounced transcriptional alterations (783 DEGs) when compared to uninfected controls with significant upregulation of Schlafen family genes. These are unknown players in pneumonia that warrant future research into their contribution to disease pathogenesis. Differential expression analysis among sickness states enabled us to identify Arginase 1 as the only predictive marker for the moribund state, which was expressed already at an early infection time-point (8hpi). This is consistent with the reported role of Arginase 1 in impairing lung-protective immunity (31). However, the detection of only a few DEGs at 8hpi is surprising given that immune cell influx especially by neutrophils is usually observed already at 6hpi (15,16). This limitation is likely due to the use of whole tissue RNA-sequencing samples, where subtle changes are masked by the complexity of the sample leading to the potential loss of small but biologically relevant signals. However, its strength lies in identifying DEGs that are either key drivers or significant consequences of infection, as they are detectable even amidst the noise of bulk sequencing.

Cell profiling of moribund mice in the acute infection phase using the ImmuCellAI-mouse tool (20), provided valuable insights into the dynamics of cell populations during severe pneumococcal infection. We found a marked reduction in B-cell populations, particularly B1 cells and plasma cells, in the lungs of moribund mice compared to uninfected controls. This depletion of B-cell subsets is interesting, as B1 cells are known to provide innate-like rapid antibody responses to polysaccharide antigens, such as the capsular polysaccharides of *S. pneumoniae* (33). The reduced presence of B- and T-cells, DCs and NK cells in the lungs was accompanied by an increase in innate immune cells, including neutrophils, monocytes, and M1-like macrophages. While these cells are essential for initial pathogen clearance, their over-activation can contribute to tissue damage and exacerbation of inflammation. For instance, excessive neutrophil infiltration is associated with alveolar damage and impaired gas exchange in pneumonia (39). The ImmuCellAI-mouse tool focuses on immune cell populations, nevertheless evaluating changes in non-immune cells, such as epithelial cells, could provide additional insights into the pathophysiology of pneumonia. Pulmonary epithelial cells play a crucial role in the recognition of pathogens, initiating immune responses through the secretion of cytokines and antimicrobial peptides (40). Tracking changes in epithelial cell activity or injury markers could reveal critical aspects of host-pathogen interactions and tissue damage during pneumococcal infection.

Comparison of DEGs from our model to those of an acute Gram-negative *Acinetobacter baumannii* pneumonia model (13) revealed substantial overlap, particularly in pathways such as IL-17, TNF, and toll-like receptor signaling. These shared responses underscore conserved innate immune mechanisms in severe bacterial pneumonia, irrespective of pathogen type. Additionally, several DEGs in our moribund mice aligned with plasma proteins associated with the Sequential Organ Failure Assessment (SOFA) score in critically ill human pneumonia patients (14), including CCL8 and CD14. These findings suggest potential translational relevance, providing a foundation for identifying biomarkers or therapeutic targets for severe pneumonia.

## Author Contributions

Conceptualization and Supervision: ADG, SW, SK

Investigation (experimental): ADG, KL, AH, IM, RG, RM, BD

Investigation (bioinformatics): ADG, JM, SW

Funding acquisition: SK, SW

Writing – original draft: ADG, SW

Writing – review & editing: all authors

## Acknowledgments

We thank the Biomedical Sequencing Facility (CeMM and Medical University of Vienna), especially Michael Schuster and Thomas Winkler-Penz, and the staff of the Core Facility of Laboratory Animal Breeding and Husbandry (Medical University of Vienna) for their support throughout the project.

## Supplementary Material

Table S1: DEGs at 2 days post-infection compared to control.

Table S2: DEGs at 30 days post-infection compared to control.

Table S3: DEGs in sick mice at 2- and 3 days post-infection compared to control.

Table S4: DEGs in moribund mice at 2- and 3 days post-infection compared to control.

Table S5: DEGs at 8 hours post-infection compared to control.

Table S6: Overlapping DEGs between *S. pneumoniae* infected mice and DEGs in an *A. baumanii* pneumonia model.

Table S7: KEGG pathways on shared DEGs in Table S6.

## Data Availability Statement

The lung tissue RNA sequencing datasets for this study can be found in the NCBI GEO database (accession number GSE289225).

## Supplementary Figure Legends

**Figure S1.**
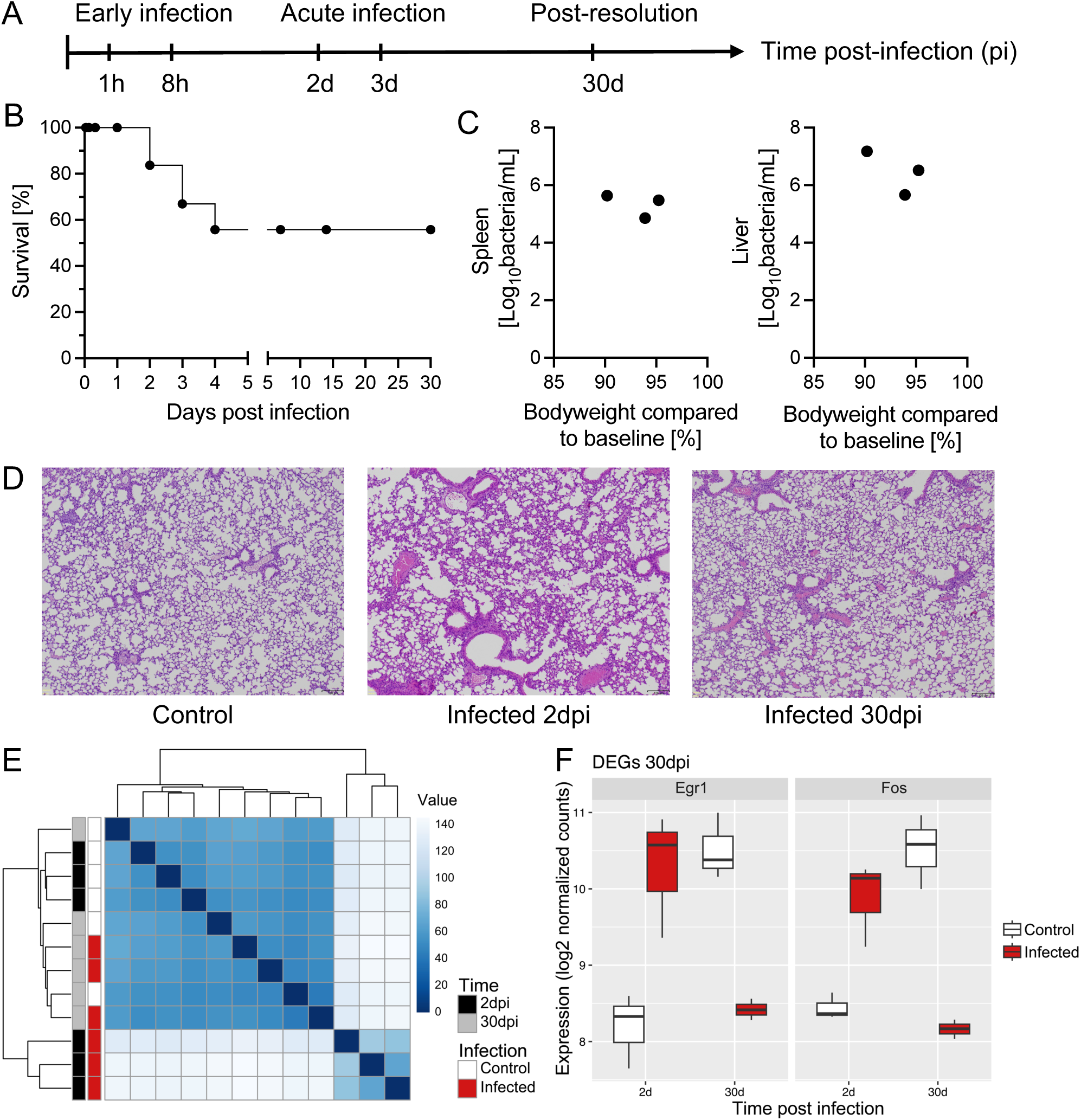
Acute and long-term alterations of pulmonary gene expression after pneumonia infection. (**A**) Female C57BL/6J WT mice were infected with 1×10^4^ CFU *S.pneumoniae* (infected) or given saline (control). Organs and blood were taken at indicated times and phases post-infection. (**B**) Survival curve of infected mice until 30dpi. (**C**) Bacterial count in spleen and liver 2dpi versus bodyweight compared to baseline. (**D**) Histology (Hematoxylin-Eosin) at 2dpi and 30dpi. (**E**) Sample distance matrix of transcriptomic lung data of infected and control mice 2dpi and 30dpi. (**F**) DEGs (absolute log2fc >2, padj <0.05, base mean count >50) 30dpi based on log2 fold change in infected versus control mice.

**Figure S2.**
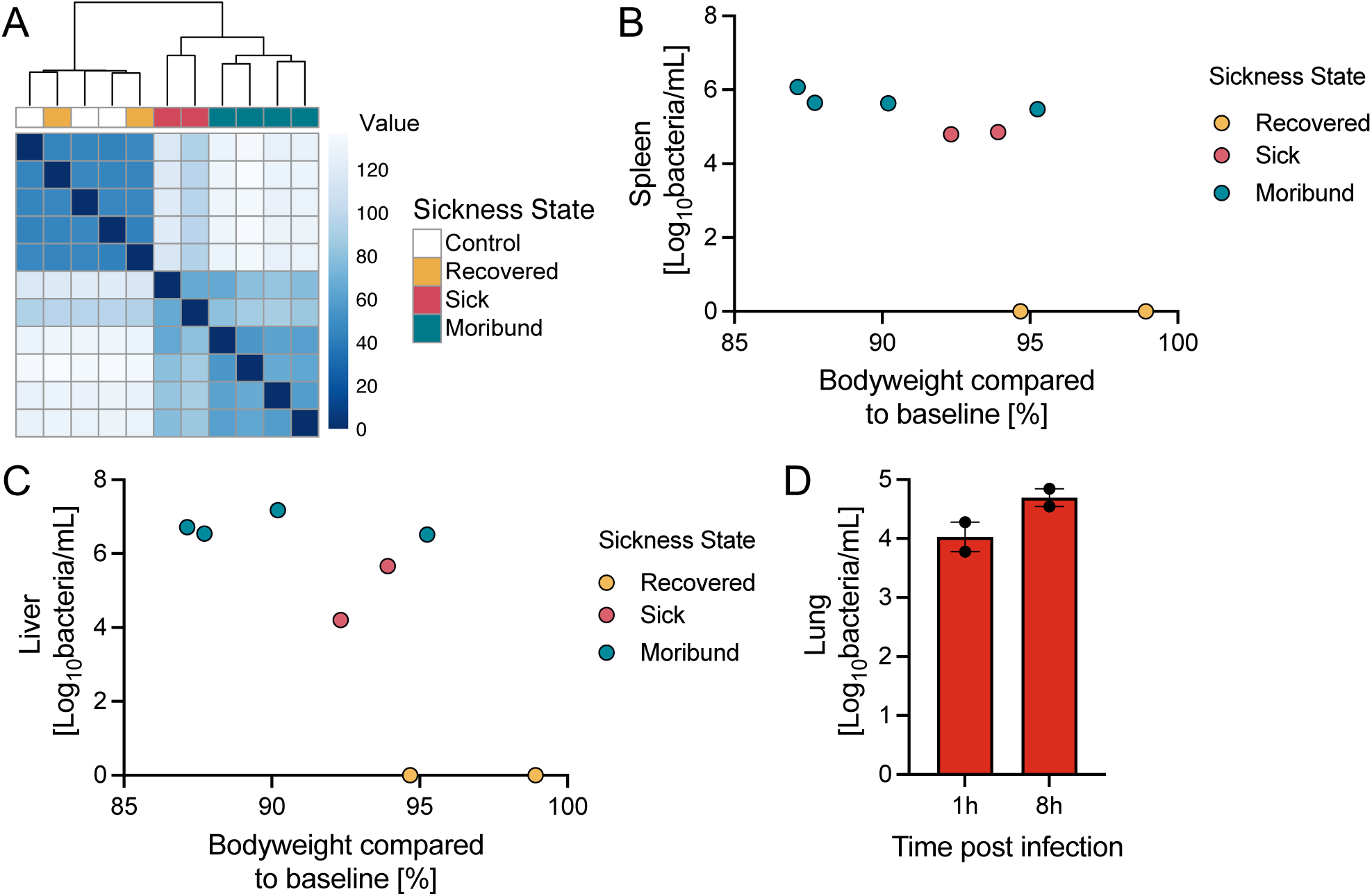
Classification of distinct sickness states during acute infection phase. Three different sickness stages (recovered, sick, moribund) were observed in *S.pneumoniae* infected mice during acute infection phase (2- and 3dpi). (**A**) Sample distance matrix of RNA seq data of infected mice (recovered, sick, moribund) and controls. (**B and C**) Bacterial count in spleen (**B**) and liver (**C**) of infected mice (recovered, sick, moribund) versus bodyweight compared to baseline. (**D**) Bacterial count in lung of infected mice 1h and 8hpi versus bodyweight compared to baseline.

**Figure S3.**
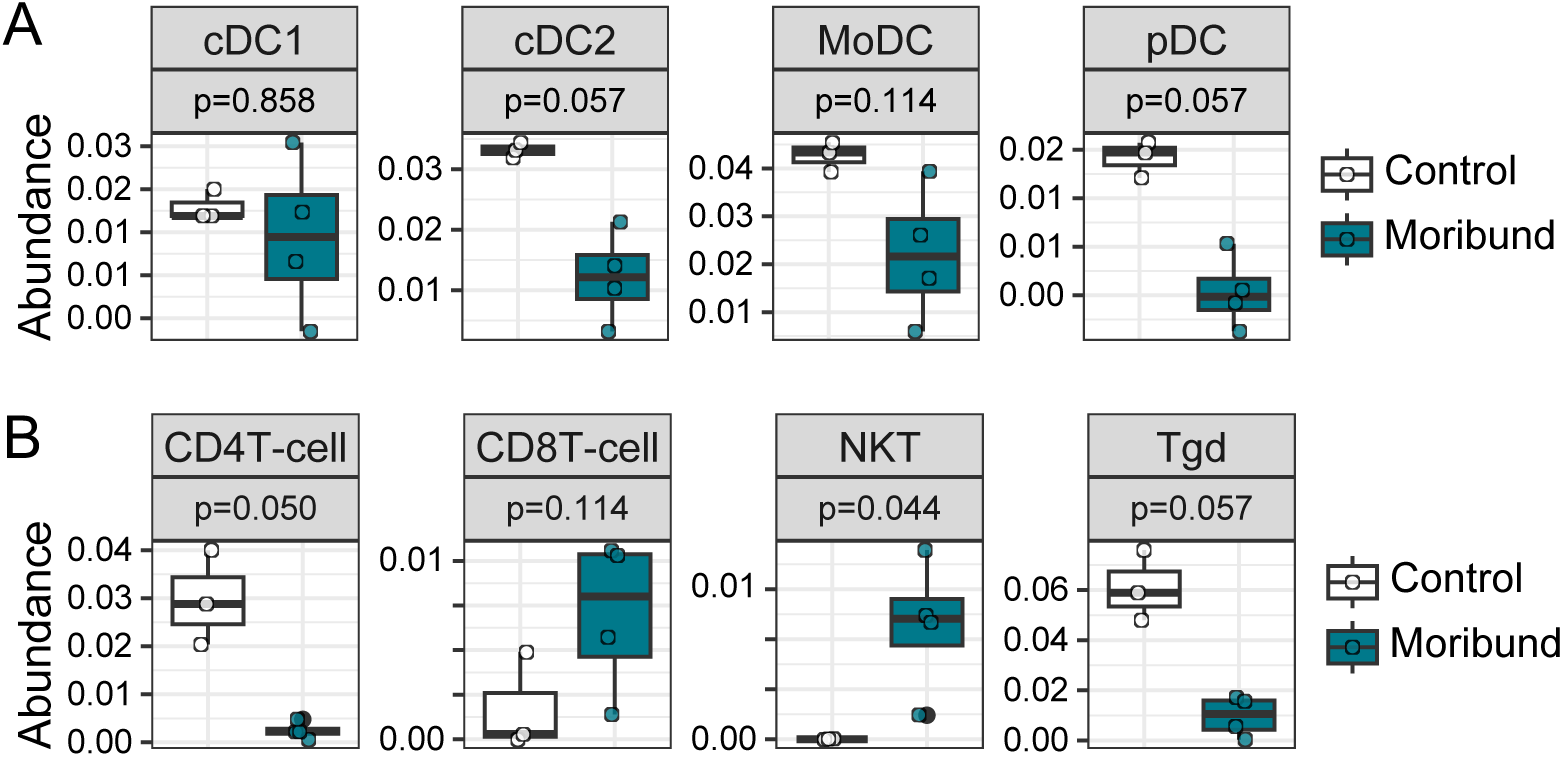
Immune cell dynamics in moribund mice during acute pneumonia. (**A to B**) Cell infiltration in lung tissue based on ImmuCellAI calculation for layer 2 dendritic cells (**A**) and T-cells (**B**) subtypes based on transcriptomic data of moribund mice and controls during acute infection phase (2- and 3dpi).

